# miniMDS: 3D structural inference from high-resolution Hi-C data

**DOI:** 10.1101/122473

**Authors:** Lila Rieber, Shaun Mahony

## Abstract

**Motivation:** Recent experiments have provided Hi-C data at resolution as high as 1 Kbp. However, 3D structural inference from high-resolution Hi-C datasets is often computationally unfeasible using existing methods.

**Results:** We have developed miniMDS, an approximation of multidimensional scaling (MDS) that partitions a Hi-C dataset, performs high-resolution MDS separately on each partition, and then reassembles the partitions using low-resolution MDS. miniMDS is faster, more accurate, and uses less memory than existing methods for inferring the human genome at high resolution (10 Kbp).

**Availability:** A Python implementation of miniMDS is available on GitHub: https://github.com/seqcode/miniMDS.

## 1 Introduction

Hi-C is a high-throughput method for genome-wide analysis of chromosome conformation. The Hi-C protocol begins by crosslinking protein-DNA interactions, with the goal of crosslinking together pairs of chromosomal loci that are proximal to one another in three-dimensional space. Crosslinked chromatin is then fragmented (typically using restriction enzymes), and the ends of the resulting fragments are marked with biotin. A random ligation reaction results either in intermolecular ligations that join two distinct DNA molecules together (also referred to as a contact) or self-ligations, which join the two ends of a single DNA molecule. After further shearing, ligation products are immunoprecipitated via the biotin label and paired-end sequenced.

The number of contacts between two loci observed in a population of cells is referred to as the contact frequency, which is inversely proportional to the average *in vivo* 3D distance between the loci in the cell population (Lieberman-Aiden et al., 2009). Contact frequency data is typically presented as a matrix, which can be analyzed using methods such as eigenvector decomposition (Imakaev et al., 2012). However, for the purposes of structural comparison and visualization, it is often useful to convert the matrix to a 3D structure or ensemble of structures.

There are two types of methods for 3D structural inference from Hi-C data (Park and Lin, 2016). Modeling-based methods assume that contact frequencies are related to distances via a probabilistic function, such as a Poisson distribution. These methods aim to maximize the likelihood of the inferred 3D structure using algorithms such as Markov chain Monte Carlo (Rousseau et al., 2011), (Park and Lin, 2016), (Varoquaux et al., 2014), (Hu et al., 2013), (Zou et al., 2016). Optimization-based methods infer a function to convert the contact frequency matrix to a distance matrix. A 3D structure is initialized (typically at random), and an objective function is used to quantify the difference between the inferred 3D structure and the distance matrix. The 3D structure is iteratively updated to minimize the objective function, using multidimensional scaling (MDS), for example (Lesne et al., 2014), (Duan et al., 2010), (Adhikari et al., 2016), (Baù and Marti-Renom, 2012), (Zhang et al., 2013), (Szalaj et al., 2016).

Recent experiments have provided Hi-C data at resolutions as high as 1-5 Kbp for several human cell lines (Rao et al., 2014). However, existing structural inference methods have not been applied to mammalian whole-genome Hi-C data with a resolution greater than 40 Kbp (Hu et al., 2013). High resolution creates unique challenges for structural inference, because computational requirements increase exponentially with the number of loci in the dataset. The sparsity of the contact matrix also increases with resolution (Park and Lin, 2016). Long-range contacts are particularly sparse, because the contact frequency between two loci decreases exponentially with the linear separation between the loci (Lieberman-Aiden et al., 2009). The excess of zeroes can be corrected using interpolation (Lesne et al., 2014) or a statistical adjustment (Park and Lin, 2016). On the other hand, zeroes provide little or no information for structural inference. Removing zeroes would thus have little informational cost and would reduce the memory and computations associated with large data sets.

We propose a structural inference method for high-resolution Hi-C data that partitions the contact matrix using a method similar to topologically associating domain (TAD) identification and performs MDS individually on each partition. This algorithm is somewhat similar to the subsampling used by other MDS approximation algorithms (Platt, 2005). Because loci preferentially interact with loci within the same TAD (Dixon et al., 2012), this approach minimizes the number of zeroes in the data. It also reduces the amount of data that must be stored in memory at any given time and allows for parallelization of analysis. The partitions are then assembled into a global structure using low-resolution data. Our method is faster, more memory-efficient, and more accurate than alternative methods and can solve a three-dimensional structure for the human genome at kilobase-resolution in less than five hours.

## 2 Methods

### 2.1 Data source

We used GM12878 Hi-C count matrices (MAPQ ≥ 30) from (Rao et al., 2014) for all analyses, which were normalized using the Knight-Ruiz normalization (Knight and Ruiz, 2013) factors provided (GEO accession number: GSE63525).

### 2.2 Algorithm

miniMDS infers detailed whole-genome 3D structures by progressively solving and integrating structures at three resolution levels. High-resolution local structures are first solved within each partition. Lower resolution structures are then inferred for each whole chromosome, and the high-resolution partitioned structures are overlaid onto these chromosomal structures after solving the optimal transformations. For inter-chromosomal analysis, a low-resolution inter-chromosomal structure is inferred, and the high-resolution intra-chromosomal structures are overlaid onto this.

#### 2.2.1 Partitioning

To solve local high-resolution structures, the contact matrix is first partitioned, with the average size of partitions determined by a user-defined smoothing parameter. Larger partitions may produce more accurate results but have greater computational requirements.

Our algorithm to partition the genome was derived from a method for identifying TADs, which used a directionality index that describes whether a locus preferentially interacts with loci upstream or downstream (Dixon et al., 2012). Loci near the beginning of a TAD preferentially interact downstream, and those near the end preferentially interact upstream. The existing method uses a hidden Markov model to identify large regions of upstream or downstream bias in Hi-C data. However, this method produces only one partitioning of the genome. To partition the genome at different length scales, we propose a modification of the directionality index method, in which directionality indices of individual loci are binned to identify large regions of bias. Partition boundaries are defined by the occurrence of a downstream-biased bin, followed by an upstream-biased bin. Partitions are then defined as the regions between boundaries. An adjustable smoothing factor determines the size of the bins (the smoothing factor multiplied by the total number of loci on a chromosome equals the bin size used in partitioning). Larger smoothing factors result in larger partitions. Given a certain smoothing factor, our method produces partitions that approximately correspond to the TADs identified by Dixon (Fig. 1). By adjusting the smoothing factor, we observed a hierarchy of genome organization (Fig. 2), suggesting that there may not be a single TAD definition for a chromosome. This is consistent with recent results demonstrating a hierarchy of genome folding, in which TADs have functional importance but no particular structural importance (Zhan et al., 2017).

**Fig. 1.**
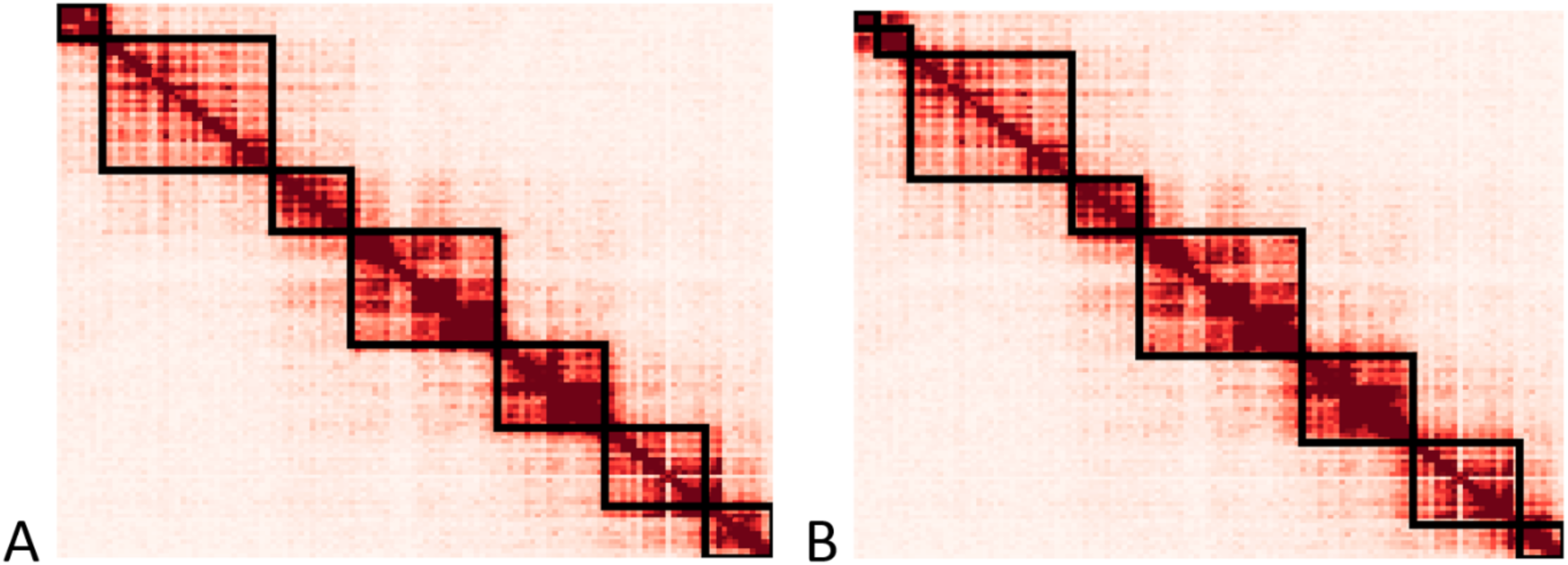
Partitions identified by the simplified algorithm with a particular smoothing factor approximately correspond to the previously identified TADs. **(A)** TADs identified by Dixon *et al*. for mouse embryonic stem cell chr6 **(B)** Partitions created by the partitioning algorithm applied to the same data.

**Fig. 2.**
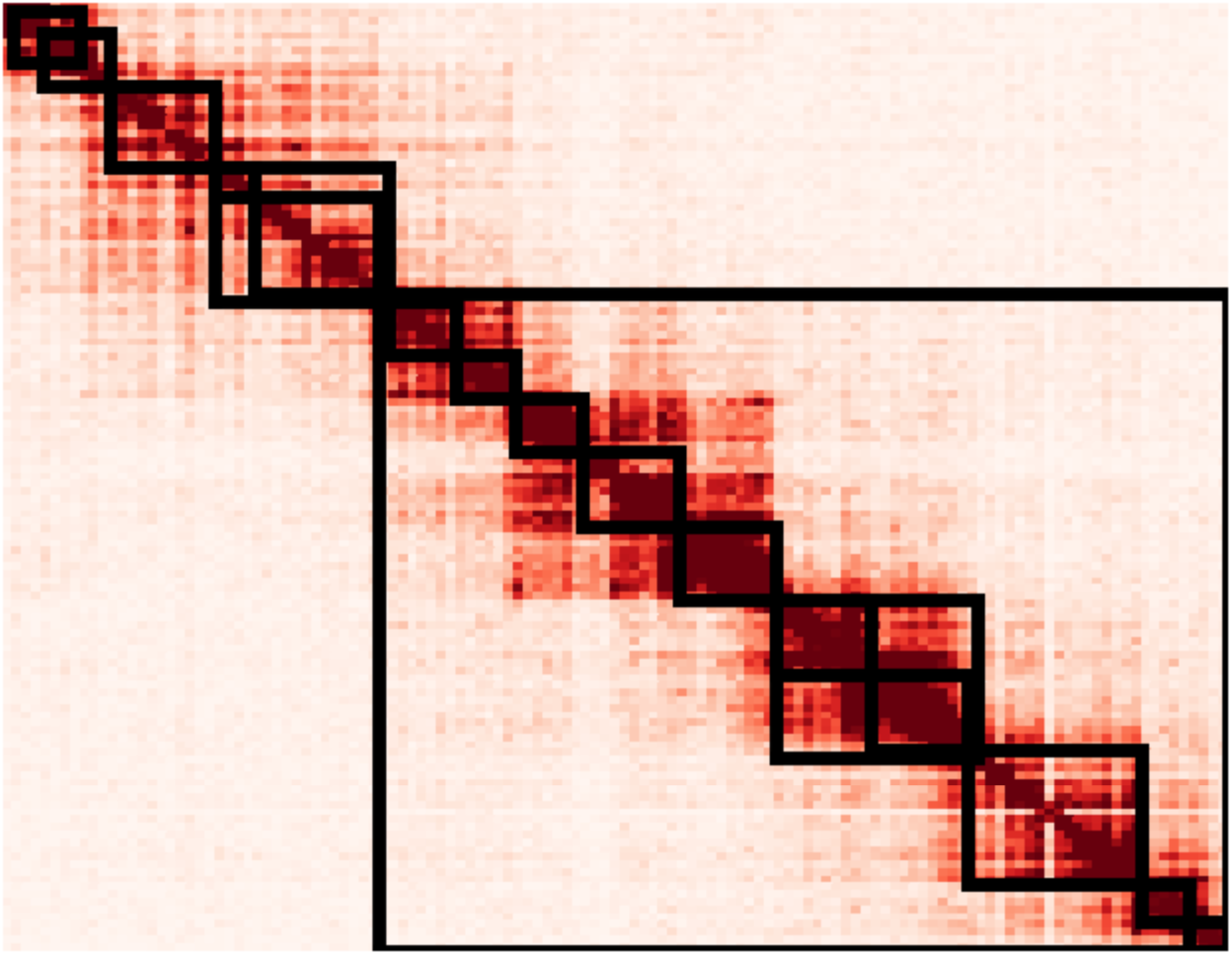
Partitions created using various smoothing factors applied to the data from Fig. 2.

#### 2.2.2 Assembling the global structure

The contact matrix for each partition is converted to a distance matrix using the following equation (zero-distances are ignored by the MDS algorithm):

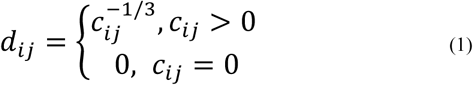
We used *α* = −4, which was found to best fit Hi-C data in results from fluorescence *in situ* hybridization (FISH) experiments (Wang et al., 2016). Inference of each partition’s structure is performed independently of every other partition, and thus is trivially parallelizable by splitting partition computations across processors. Three-dimensional metric MDS (mMDS) is performed on each partition’s distance matrix in parallel, producing a high-resolution structure for each partition (Fig. 3A). To infer the spatial relationships between partitions, a global structure is inferred from low-resolution data, using mMDS without partitioning (Fig. 3C). High-resolution partitions are then assembled in parallel, using the low-resolution structure as a guide. By taking the average coordinates of multiple loci, each high-resolution partition structure is approximated at the same resolution as the global low-resolution structure (Fig. 3B). As a result, each low-resolution locus in a partition structure has an analog in the global low-resolution structure. The optimal transformation (translation, rotation, and reflection) to align each low-resolution partition to its analog in the global structure (Fig. 3D) is calculated using the Kabsch algorithm (Kabsch, 1976). Each transformation is then applied to the corresponding high-resolution partition structure (Fig. 3F) to create a global high-resolution intra-chromosomal structure (Fig. 3G). To infer inter-chromosomal structures, a global inter-chromosomal structure is inferred at very low resolution, analogous to Fig. 3C. miniMDS is applied to each chromosome in the structure individually. Then the high-resolution miniMDS intra-chromosomal structures are aligned to the inter-chromosomal structure.

**Fig. 3.**
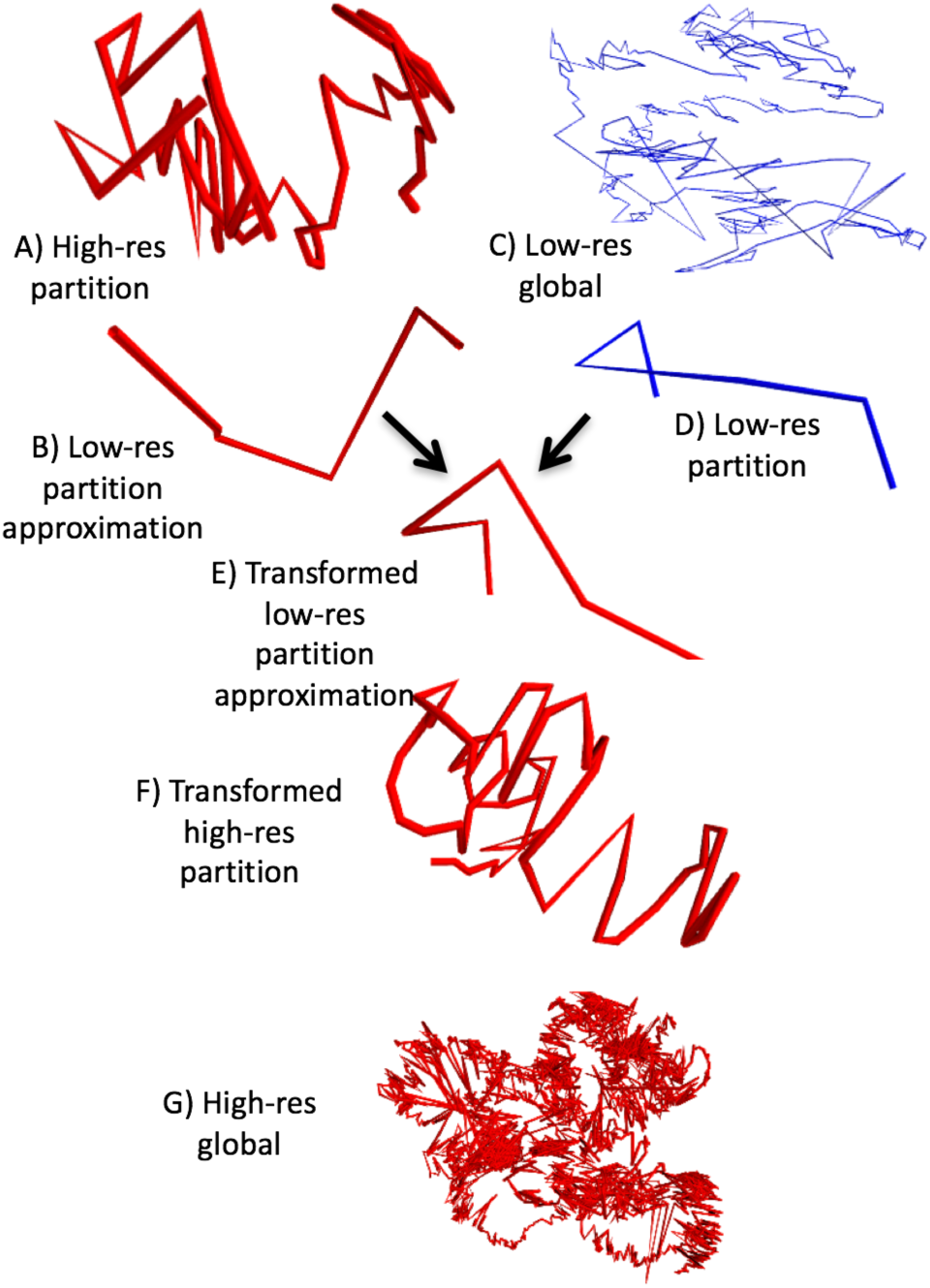
Overview of the miniMDS algorithm, demonstrated on GM12878 chr22 at 10-Kbp resolution. MDS is applied to each partition individually, creating high-resolution local structures,such as the structure shown in **(A)**. The structure is approximated at low-resolution, **(B)**. MDS is performed on a low-resolution dataset, creating a global low-resolution structure, **(C)**. The low-resolution partition has an analog in the global structure, **(D)**. The optimal transformation to align the low-resolution approximation with its analog is calculated. For illustration, **(E)** shows the transformation applied to the low-resolution approximation. The transformation is applied to the high-resolution partition, **(F)**. When this process is repeated for all partitions, a global high-resolution structure is created, **(G)**.

### 2.3 Implementation

The miniMDS algorithm is implemented in Python. MDS steps are performed using scikit-learn with default parameters (Pedregosa et al., 2011). Parallelization was performed using pymp. miniMDS is released under an MIT open source license. miniMDS source code is available on GitHub: https://github.com/seqcode/miniMDS.

### 2.4 Comparison with alternative approaches

We attempted to test the performance of BACH (Hu et al., 2013), MOGEN (Trieu and Cheng, 2014), 3D-GNOME (Szalaj et al., 2016), ChromSDE (Zhang et al., 2013), tREX (Park and Lin, 2016), MCMC5 (Rousseau et al., 2011), PASTIS (Varoquaux et al., 2014), TAD-bit (Baù and Marti-Renom, 2012), HSA (Zou et al., 2016), Chromosome3D (Adhikari et al., 2016), and InfMod3DGen (Wang et al., 2015) (Table 1). We also tested classical MDS (cMDS), which is used by ShRec3D (Lesne et al., 2014), and standard mMDS.

**Table 1.**
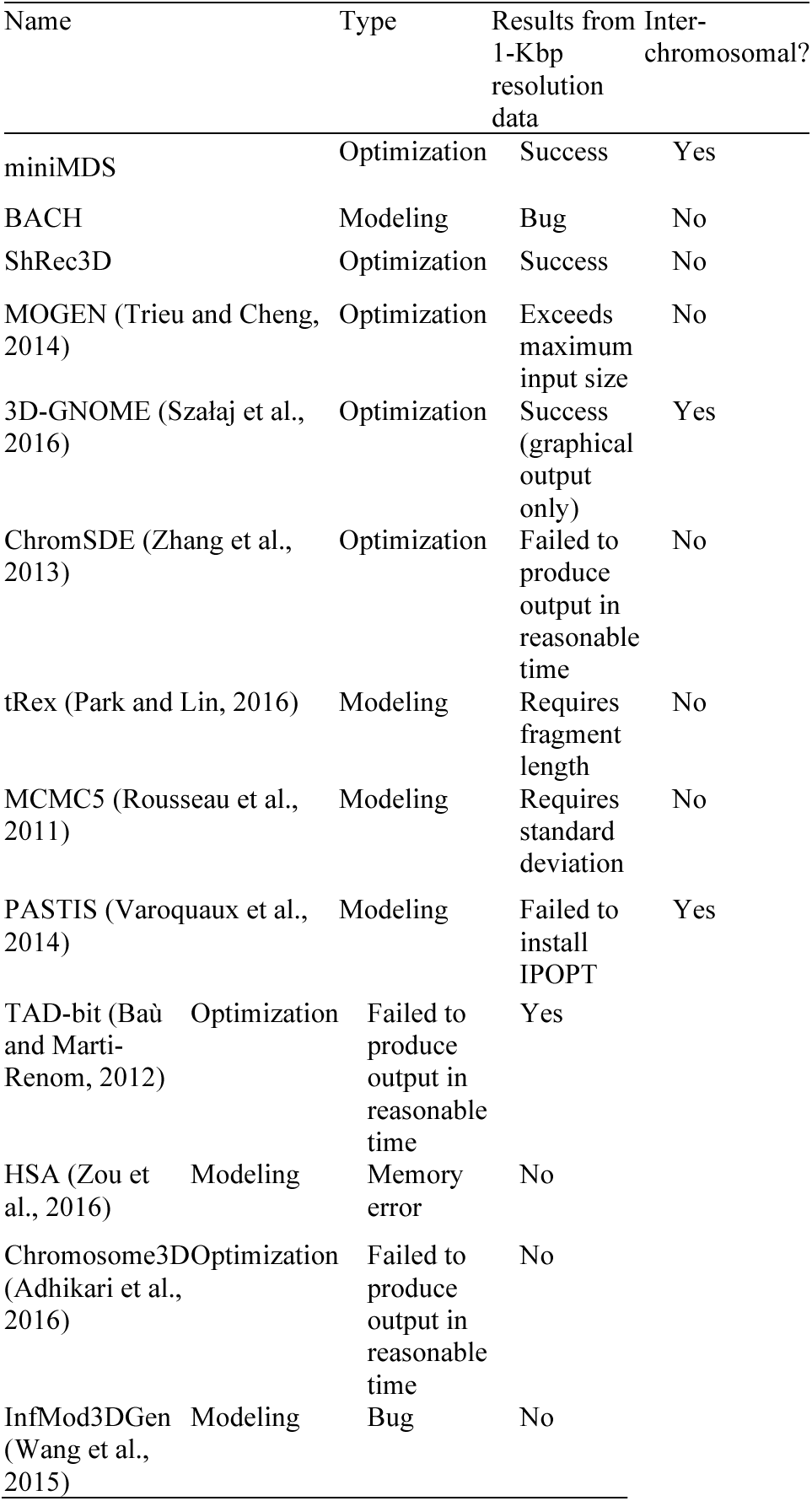
Comparison of Hi-C structural inference methods

We were unable to install PASTIS, because it requires the IPOPT package, which must be built from source and has many additional dependencies. We were unable to test MCMC5 because it requires the standard deviation of contact frequencies, which was not available for the datasets we used. We were unable to test tREX because it requires fragment length, which was not available for the datasets we used. PASTIS, MCMC5, and tREX are modeling-based methods. Because modeling-based methods must probabilistically explore parameter space, they have large computational requirements. Thus it is unlikely that they would have been able to analyze high-resolution data if we had been able to run them. BACH and InfMod3DGen were excluded from further testing due to errors. BACH produced an error for chr22 100-Kbp-resolution data. InfMod3DGen produced an error when tested on its sample data for yeast chrXVI at 100-Kbp resolution. A quadratic solver was used for ChromSDE. Defaults were used for all other parameters.

Table 1 summarizes the results of Hi-C structural inference methods applied to chr22 kilobase-resolution data. The only algorithms that were able to infer structures at kilobase-resolution were miniMDS, 3D-GNOME, and ShrRec3D. We were unable to evaluate the accuracy of 3D-GNOME because it provides only graphical output, rather than 3D coordinates, so it was excluded from further testing. ShRec3D is represented in further testing by cMDS.

## 3 Results

### 3.1 miniMDS is more efficient than other structural inference methods

We tested the computational time required for miniMDS, standard mMDS, cMDS, TAD-bit, MOGEN, Chromosome3D, and ChromSDE to analyze chr22 Hi-C data at 10-Kbp resolution (Fig. 4) and 100-Kbp resolution (Supplemental Fig. 1). TAD-bit did not produce output in a reasonable amount of time (i.e. several days) and thus was excluded from further analysis. HSA was excluded from the 10-Kbp analysis because of the large amount of time required for the 100-Kbp resolution (over 60 hours), which we assumed would be significantly greater for 10-Kbp resolution. miniMDS had the lowest time requirements of all remaining methods at both resolutions.

We also tested the amount of RAM required for miniMDS, standard mMDS, cMDS, MOGEN, Chromosome3D, and ChromSDE to analyze chr22 data at 100-Kbp resolution (Supplemental Fig. 2). Though the full benefits of miniMDS are not demonstrated at low resolution, miniMDS had the lowest memory requirements. MOGEN, Chromosome3D, and ChromSDE required orders of magnitude more memory.

**Fig. 4.**
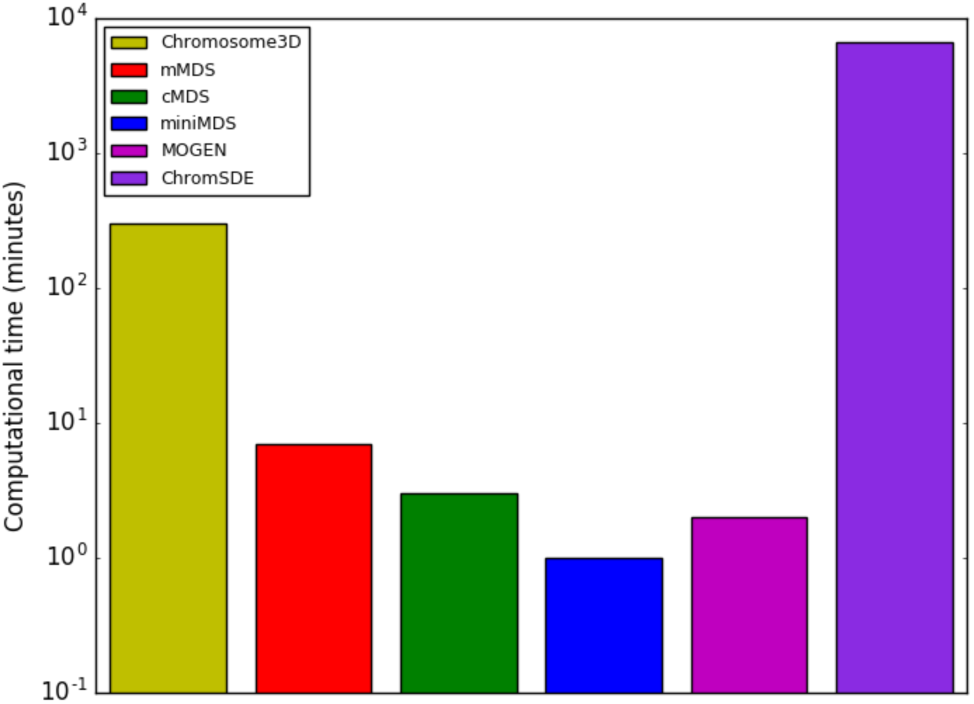
Time required to analyze chr22 data at 10-Kbp resolution

### 3.2 miniMDS is robust

We performed two iterations of Chromosome3D, mMDS, miniMDS, MOGEN, HSA, and ChromSDE on chr22 at 100-Kbp resolution. cMDS was excluded because it does not use a random seed, so it produces the same output from every iteration. We created a distance matrix from each set of output coordinates and calculated the Pearson correlation between the distance matrices for each pair of iterations (Fig. 5). MDS-based methods, Chromosome3D and ChromSDE produced the same output from both iterations, resulting in correlations of 1. The correlation between different iterations was lower for HSA and MOGEN.

**Fig. 5.**
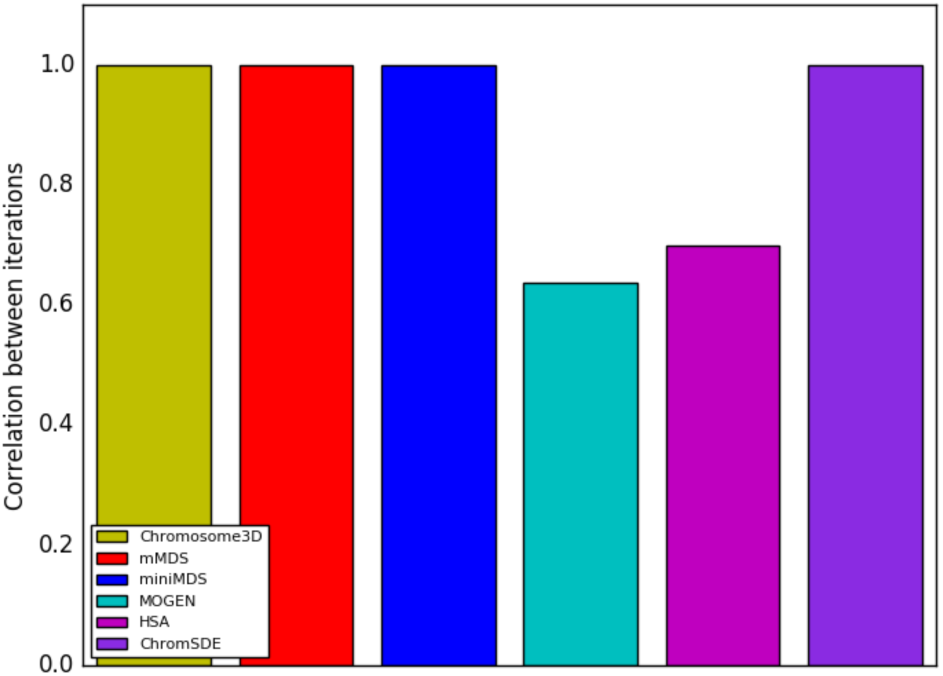
Correlation between the output of two iterations of the same method applied to the same chr22 100-Kbp-resolution dataset

### 3.3 Performance on 10-Kbp-resolution data

#### 3.3.1 Time and memory requirements

To determine how computational requirements changed with the number of loci, we tested MOGEN, miniMDS, mMDS, and cMDS on all chromosomes at 10-Kbp resolution. Chromosome3D, ChromSDE, and HSA were excluded from these analyses because the time costs were prohibitive. MOGEN was the fastest on average, with miniMDS performing almost as well (Fig. 6). However, given that MOGEN is not robust (see section 3.2), its speed may be because the algorithm does not converge. Both methods were significantly faster than standard mMDS and cMDS, with a lower rate of increase of time requirements with the number of loci. miniMDS had the lowest memory requirements, with minimal increase in memory requirements for increasing number of loci (Fig. 7).

**Fig. 6.**
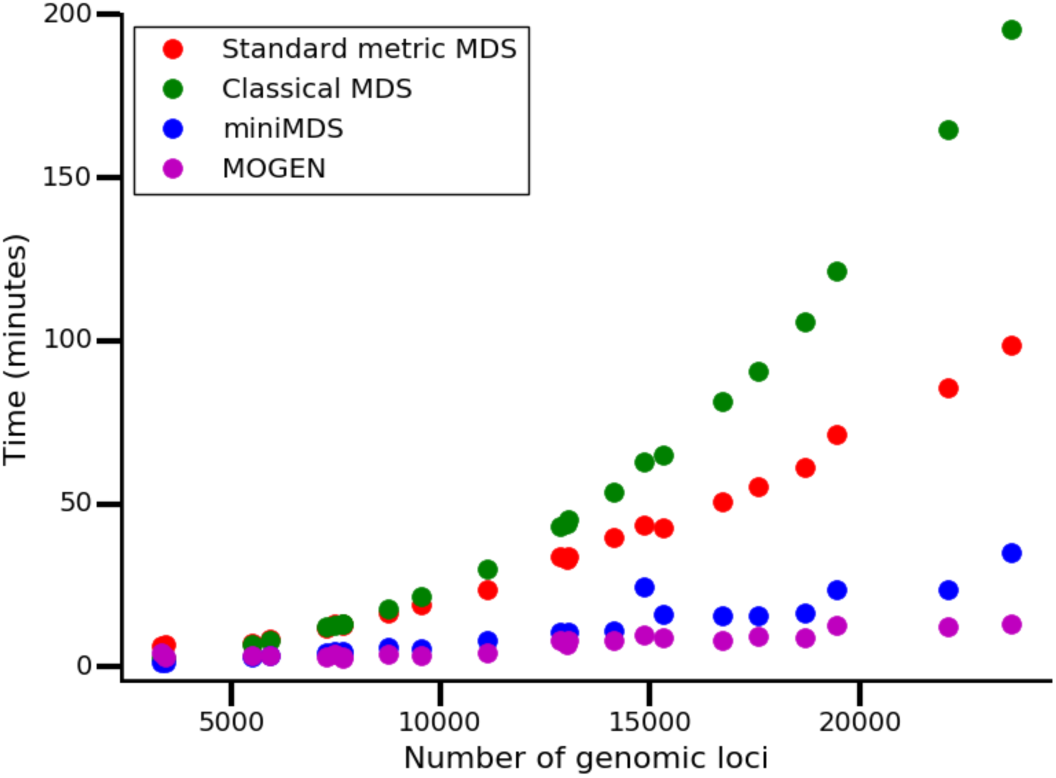
Time required to analyze each chromosome at 10-Kbp resolution

**Fig. 7.**
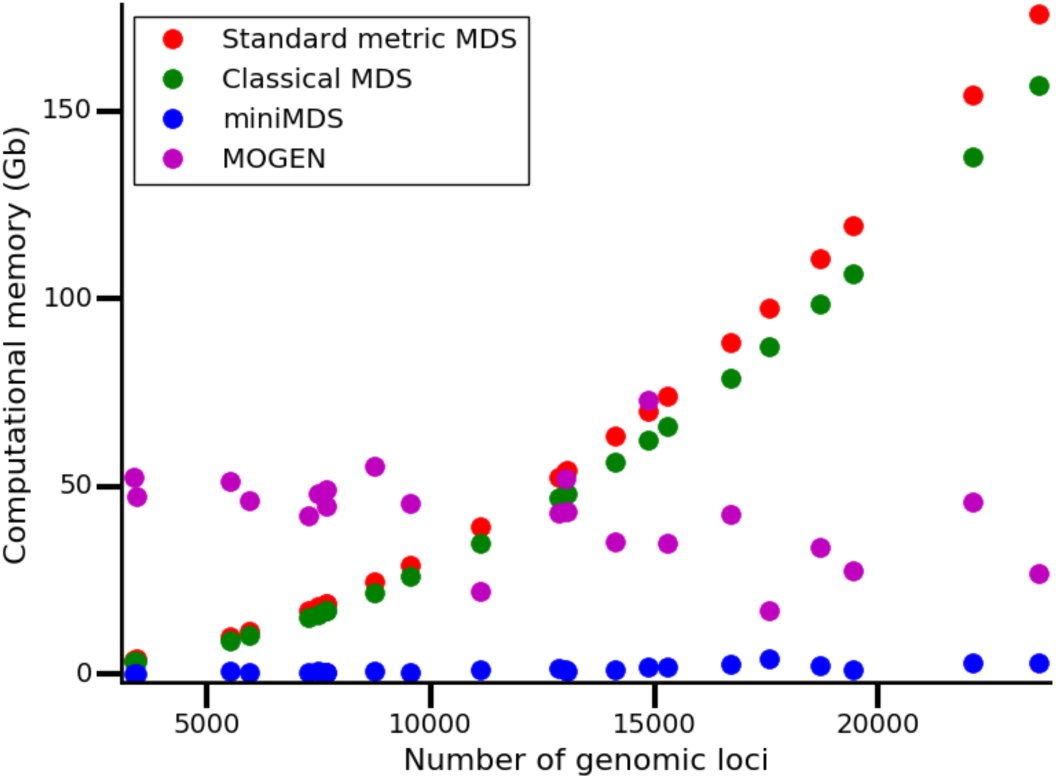
Memory required to analyze each chromosome at 10-Kbp resolution

#### 3.3.2 Correlation between input and output

We calculated the correlation between input distances and distances inferred from the output 3D structure for MOGEN, miniMDS, mMDS, and cMDS applied to each chromosome at 10-Kbp resolution. For the MDS methods, accuracy was calculated as the correlation between the input distance matrix and the distances calculated from output coordinates. For MOGEN, we inferred *α* by fitting an exponential curve to distances, which were calculated from output coordinates, as a function of contact frequencies. We then transformed contact frequencies to inferred distances using the inferred value of *α* and calculated the correlation between inferred distances and output distances.

The correlation was highest for miniMDS for every chromosome (Fig. 8), demonstrating that miniMDS infers 3D structures that are more consistent with the underlying Hi-C data. The structures inferred for chr22 using mMDS and cMDS are collapsed globules with few discernable features (Fig. 9A-B). MOGEN produces a fairly unfolded structure dominated by outliers, with little evidence of TAD structure or the hierarchical organization hypothesized by the fractal globule model ((Rao et al., 2014) (Fig. 9D). miniMDS produces a structure that is densely clustered but still displays discernable structures (Fig. 9C).

MOGEN and MDS-based methods are optimization-based. We were interested in whether a modeling-based method would offer advantages for accuracy. HSA was the only modeling-based method that we were able to test (see section 2.4), so we evaluated its performance on 100-Kbp-resolution chr22 data, even though it is not robust (see section 3.2). We calculated its correlation between input and output using the analysis described for MOGEN. The correlation was lower for HSA than for MDS-based methods applied to the same data (Supplemental Fig. 3).

**Fig. 8.**
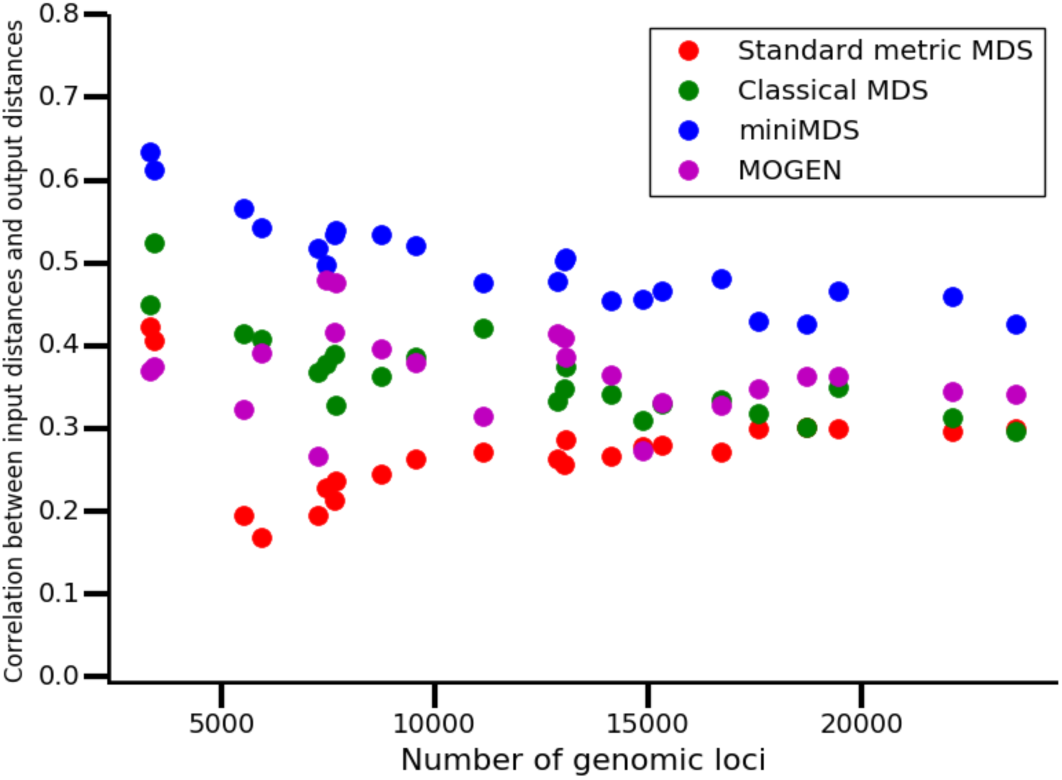
Accuracy of methods for each chromosome at 10-Kbp resolution

**Fig. 9.**
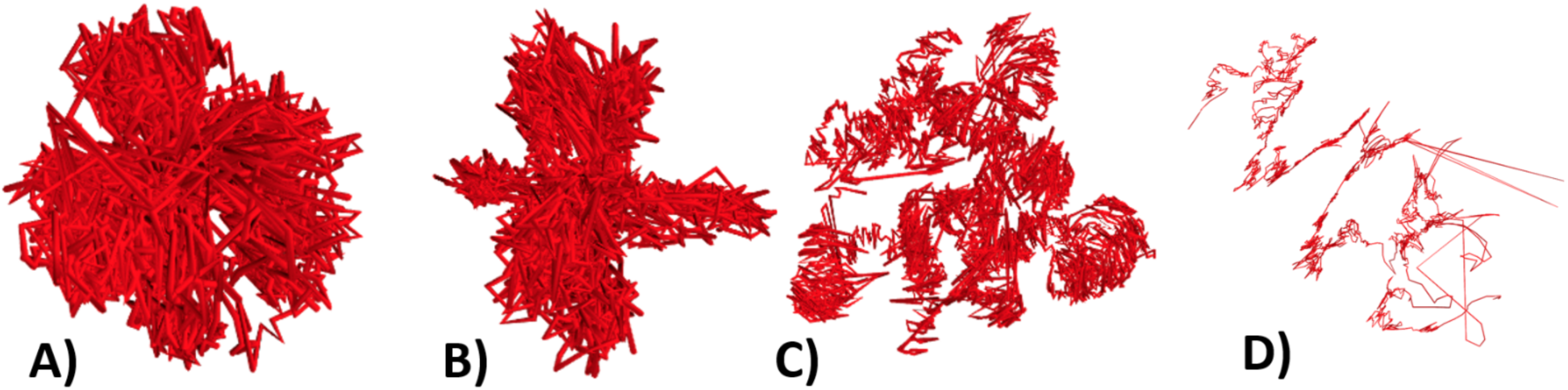
Chr22 10-Kbp-resolution structures produced by mMDS (A), cMDS (B), miniMDS (C), and MOGEN (D).

### 3.4 miniMDS performs whole-genome structural inference at high resolution

We used miniMDS to infer a global conformation for all GM12878 chromosomes using 10-Kbp-resolution Hi-C data (Rao et al., 2014) (Fig. 10). We used a smoothing factor of 0.05 and a minimum partition size of 1% of the matrix. Intermediate low-resolution structures for each whole chromosome were inferred from 100-Kbp-resolution data. Inference of the entire whole-genome structure required 4 hours 21 minutes.

**Fig. 10.**
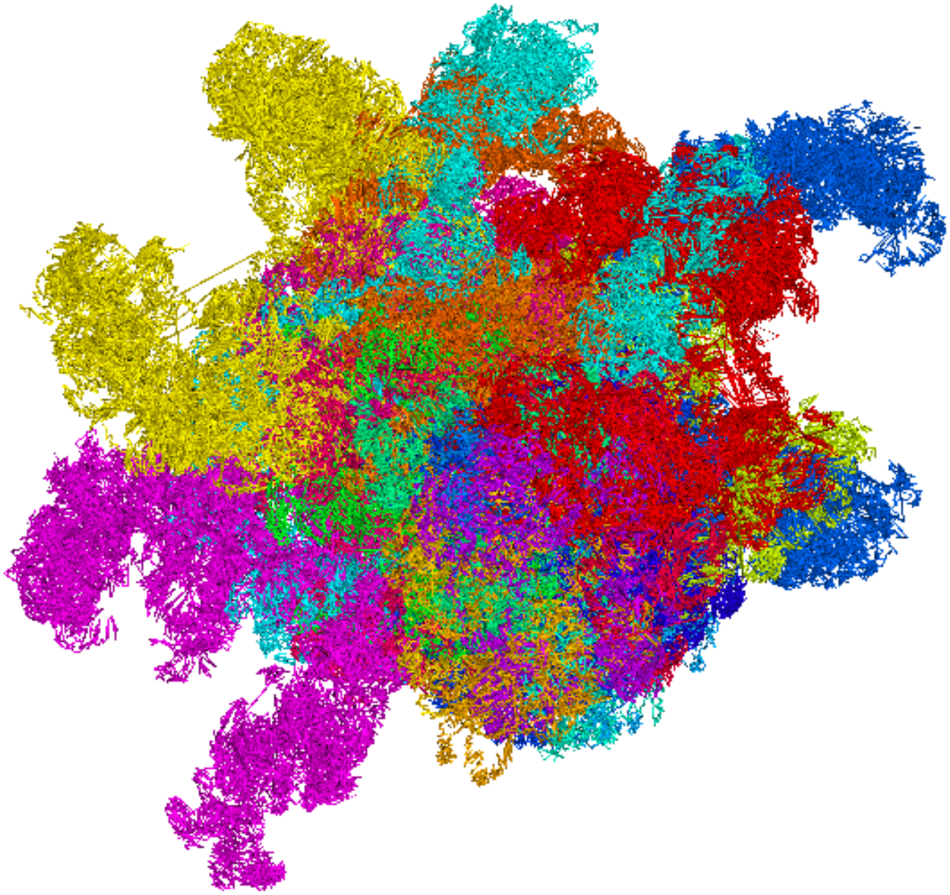
GM12878 chromosome conformation at 10-Kbp resolution

## 4 Discussion

We have presented miniMDS, a method for inferring 3D structures from Hi-C experiments that is suitable for high-resolution data. It uses genome partitioning and parallelization to achieve greater speed and lower memory requirements compared to alternative methods. Theoretically, the memory requirements of miniMDS could always be maintained below a certain threshold by increasing the number of partitions, regardless of the size of the dataset. As measured by the correlation between input distances and output distances, miniMDS is more accurate than other methods that are able to efficiently analyze large, high-resolution datasets. However, we note that Chromosome3D and ChromSDE, which do not scale well to large datasets, perform better on this metric for smaller datasets. These algorithms use different optimization algorithms, which could be made more efficient using a partitioning strategy similar to miniMDS.

Comprehensive identification of chromatin loops, which play an important role in gene regulation, requires high-resolution Hi-C data (Rao et al., 2014). Detection of co-localization of DNA-binding proteins would also be improved by high resolution. Though Hi-C data can be analyzed in two dimensions, the dynamics of chromosome conformation are easiest to understand in three dimensions. For example, 3D structures from multiple time points or cell types could be compared using a structural alignment algorithm (Hasegawa and Holm, 2009) to determine how the localization of genomic regions changes. It is possible that dynamic genomic regions are correlated with changes in gene regulation. High-resolution 3D chromosome conformation inference will thus contribute to exploration of the “4D nucleome” (Chen et al., 2015). miniMDS is the only method currently available that can accurately solve whole-genome 3D structures from the highest-resolution Hi-C datasets.

## Acknowledgements

The authors thank the members of the Center for Eukaryotic Gene Regulation at Penn State for helpful feedback and discussions.

